# Simplified and unified access to cancer proteogenomic data

**DOI:** 10.1101/2020.11.16.385427

**Authors:** Caleb M. Lindgren, David W. Adams, Benjamin Kimball, Hannah Boekweg, Sadie Tayler, Samuel L. Pugh, Samuel H. Payne

## Abstract

Comprehensive cancer datasets recently generated by the Clinical Proteomic Tumor Analysis Consortium (CPTAC) offer great potential for advancing our understanding of how to combat cancer. These datasets include DNA, RNA, protein, and clinical characterization for tumor and normal samples from large cohorts in many different cancer types. The raw data are publicly available at various Cancer Research Data Commons. However, widespread re-use of these datasets is also facilitated by easy access to the processed quantitative data tables. We have created a Python package, `cptac`, which is a data API that distributes the finalized processed CPTAC datasets in a consistent, up-to-date format. This consistency makes it easy to integrate the data with common graphing, statistical, and machine learning packages for advanced analysis. Additionally, consistent formatting across all cancer types promotes the investigation of pan-cancer trends. The data API structure of directly streaming data within a programming environment enhances reproducibility. Finally, with the accompanying tutorials, this package provides a novel resource for cancer research education.

## Introduction

Large consortia, like the Clinical Proteomic Tumor Analysis Consortium (CPTAC), drive science forward by creating coordinated and structured datasets on a scale that is typically not possible with individual investigators. They amass both a number of samples and diversity of measurements that requires a large collaborative effort. In addition to the primary analysis done by the consortium and published as flagship manuscripts^1–6^, these datasets are designed to be a resource to the scientific community to explore new questions or apply novel methodologies^7–11^.

Proteogenomic cancer data is of interest to a wide interdisciplinary audience and different scientists may want to interact with different data products, e.g. raw instrument data versus summarized quantitative tables. Funding agencies have focused building resources for the dissemination of voluminous raw data files, e.g NCI Genomic Data Commons^12,13^. These data warehouses address the logistical and technological challenges of storing and disseminating terabytes of sequencing and mass spectrometry data. However, raw data repositories have a limited audience, as the re-analysis of raw instrument files requires domain-specific knowledge and significant computational resources.

A growing audience of scientists want to directly interact with quantitative proteogenomics data tables and not the raw data. Currently there is not a common method for projects, large or small, to share these data tables in an open and computable format. Although some quantitative data may be shared through the large data warehouses^14^, this mechanism has several drawbacks. First, the final data tables used in a publication are usually highly curated and processed by harmonization, batch correction, normalization, filtering, etc. The detailed attention in these computational adjustments should not be lost; the public should have access to the exact data tables used in a publication. Second, the Data Commons model as currently designed separates multi-omics datasets into different Data Commons instances, requiring users to have prior knowledge of which datasets belong together and where they are stored. Finally, computational convenience should be a driving factor in the data storage mechanism, meaning that data should be easy to access programmatically and to quickly use in computation. Thus, an alternative dissemination mechanism that facilitates accessing and utilizing these data tables is needed to serve a broader scientific community. A convenient method for disseminating coordinated datasets is the data API model, where data is streamed directly into a programming environment^15^.

As an illustration of the need for a higher-level data distribution methods, consider the widespread use of Jupyter Notebooks^16,17^ and other similar programming environments^18^ for sharing research. These tools are ideal for both explaining the context of an analysis, and showing the exact methodology. However, for this method of sharing research to be successful, it needs an accompanying flexible data distribution method. If a shared notebook uses files that are stored on the original researcher’s computer or are too large to be conveniently streamed, it will be much less useful to others who wish to replicate and extend that analysis. The data API model solves this problem by ensuring that the exact version of a dataset is globally and universally accessible.

We present here a data API for proteogenomic cancer data generated by CPTAC, representing six tumor types. All tumor samples are characterized with genomic, transcriptomic, proteomic and clinical data. The API streams proteogenomic data directly into a Pandas dataframe within a Python programming environment, dramatically improving the simplicity of data access. By using native dataframe variables, the proteogenomic data easily integrates with common Python libraries for machine learning, graphing and statistics. Along with the API, we released an extensive set of tutorials to demonstrate common data analysis methods in proteogenomic and pan-cancer studies.

## Methods

### Overview

When reading these methods, it is important to keep in mind the distinction between the data files and the software API used to access them. In our project, these two entities are entirely separate and can be changed/updated independently.

The CPTAC Python package, called `cptac`, is a data API that facilitates access and utilization of cancer proteogenomic data within Python scripts (Figure 1). Similar to other Data-as-a-Service applications, the `cptac` package gives users on-demand access to structured data. The API parses, loads, integrates and manipulates the cancer proteogenomic data from the CPTAC consortium. When data are accessed via the API, they are presented to the user as Pandas dataframes. This data structure conveniently and seamlessly integrates with most major machine learning, graphing and data analytic libraries in Python.

**Figure 1.**
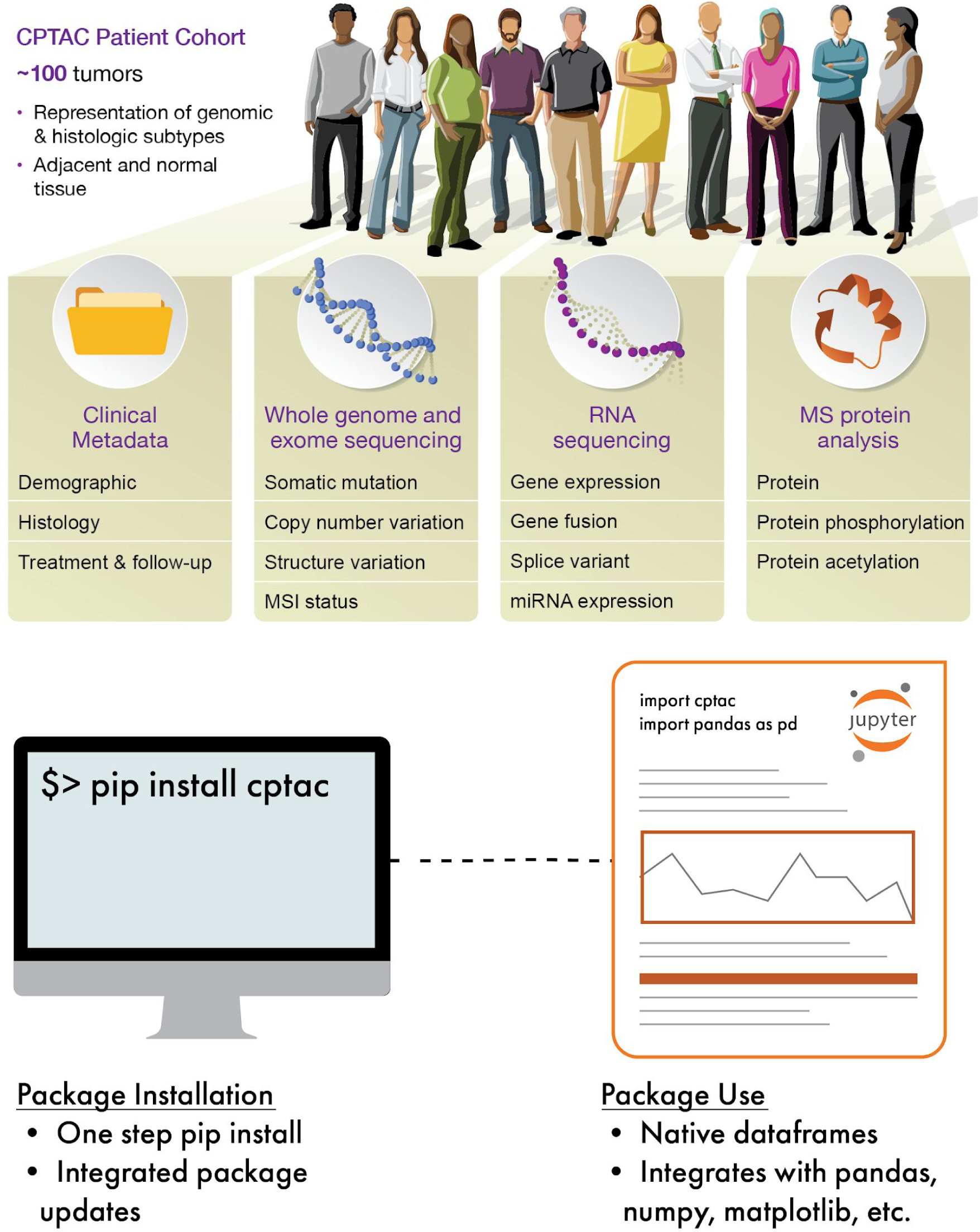
CPTAC data API. (top) Cohorts in CPTAC have the same fundamental multi-omics and clinical data. (bottom) The `cptac` Python module is simple to install and use within a Python programming environment.

All the software code for the `cptac` package is open source and available at (https://github.com/PayneLab/cptac). Formal versions are tagged on GitHub and released as software updates through the Python Package Index (https://pypi.org/project/cptac/). Thus, **developers** can work with the code of the API by forking the GitHub repository, but **users** of the data should install the package on their local computer with “`pip install cptac`”. Users should follow the tutorials at ∼/docs for explicit code demonstrating the use of the API. Developers should follow the instructions in ∼/devdocs for software design and implementation requirements.

### Data Tables

The data accessible with the `cptac` package are the published final data tables for each CPTAC tumor type, namely the published data tables associated with flagship publications of the CPTAC consortium for six cancer types: colon^5^, serous ovarian^4^, breast [*in press*], endometrial^2^, renal clear cell^1^, and lung adenocarcinoma^3^. The dataset for every tumor type contains genomic, transcriptomic, proteomic and phosphoproteomic measurements on the tumor samples. Each patient in the dataset is described by a variety of clinical data including demographic, treatment and outcome.

### Package Construction and Organization

The methods of tumor collection and data generation within CPTAC follow a consistent, organized structure (Figure 1A). Our package has taken advantage of this consistency by creating an abstract class ‘Dataset’ which defines common variables and ‘get’ functions to access all data tables that a dataset will have, e.g. get_clinical(), get_CNV(), get_proteomics(), get_somatic_mutation(), etc. In addition to simple data access, the `cptac` package assists users in merging data across dataframes using native *join* functions as part of the abstract Dataset class. These work on any combination of omics data and metadata, e.g. join_metadata_to_mutations(). The *join* functions facilitate all merging to ensure that the returned dataframes are properly keyed and indexed. Each tumor type is coded as a class object that inherits from the abstract Dataset class, i.e. a derived class. Thus, each derived class only needs to parse its specific data files into Pandas dataframes matching the format required by the abstract Dataset container, and then the rest of the functionality is automatically integrated. This construction greatly simplifies the process of adding new tumor types.

To minimize the number of dataframes and keep data descriptors present in the same variables as data values, we have implemented a multi-level index for some omics dataframes. For example, proteomics and transcriptomics dataframes reference protein/RNA isoforms both by their common name and by their unique database identifier. Although common names are used as the primary indexing key, the unique database identifiers are used as the secondary key to differentiate between isoforms. A second data type which uses a multi-index is PTM proteomics data such as phosphorylation. Here the multi-index comprises gene name, database identifier, modified amino acid residue(s) and also the MS-identified peptide sequence. This is necessary because multiple peptides may be observed in a dataset with the same phosphorylated residue, often arising because of incomplete tryptic digestion.

Finally, the `cptac` package contains extra functionality within the `utils` sub-package, accessed as “`from cptac import utils`” or “`import cptac.utils`”. The `utils` sub-package implements commonly used functions and is continually expanding. To help users identify interacting proteins, a set of utility functions access pathway information from BioPlex, STRING, Uniprot and WikiPathways. The `utils` also provides several wrappers for common statistical tests, like t-tests and linear regression, with automatic correction of the p-value cutoff for multiple hypothesis testing. Finally, `utils` automates the identification of frequently mutated genes.

### Streaming data through the API

The `cptac` module gives users access to a large amount of data; the disk space for a single cancer type is 50 - 100 MB. Because PyPI restricts the package size, and because a user may not want all the data for all cancer types, the data files are not stored directly within the package or its GitHub repository. Initially, the package contains only the URLs needed to access the data. The user must request to download the dataset for a particular cancer type, whereon the package uses one of these URLs to download an index file for that dataset. This index file contains a list of all versions of the dataset, and a list of files, URLs and MD5 hashes contained in each version. After each file is downloaded, the package hashes it and checks against the hash in the index, to make sure none of the data was corrupted in the download process. After a user has downloaded a dataset, they can load the data into variables in their Python program, e.g. “`dataset = cptac.Colon()`”. The current implementation of the `cptac` package utilizes Box as a remote storage server. However, the software architecture has isolated the code involving remote data streaming, which makes it trivial to change the remote location.

### New data or API releases

The CPTAC data changes periodically during analysis as various pipelines are compared and optimized. This is tracked by the consortium as versioned data releases, e.g. 1.0 or 2.0, etc. The package has access to formal release versions and automatically monitors whether a user is working with the most recent release. Each time the user loads a dataset within their Python code, the package checks to make sure that the user has the current index for that dataset. If necessary, it downloads a new version of the index from the server. Then, it checks whether the user is using the latest version recorded in the index. If the user didn’t request the latest version, the package warns them that they are using an out-of-date version. It then loads whichever data version the user requested.

Note that when new data versions are released and downloaded, the package maintains any prior versions that were downloaded to a user’s computer. When a user loads a dataset, they can tell the package to load one of these older versions. Thus, if changes in a new version affect the results of a user’s analyses, they can go back and look at the old version to compare the two outputs explicitly.

The package also makes sure that the software API itself is up to date. Periodically we release new versions of the package to add or improve functionality. Each time the user imports the package, it downloads a small file from the server that contains the latest release version number. It then compares this with the version number of the installed copy of the package. If they don’t match, it informs the user that their copy of the package is out-of-date, and tells them how to update it using `pip`.

As internet connectivity is not universal, the package first checks to see whether there is a connection prior to comparing index files for API or data versions. If it is not possible to download the files needed to check whether the indices and package are up-to-date, the package automatically skips this version check and works with whatever it has installed locally.

### Error Handling

The table joining and other data manipulation functions in our package provide novel opportunities for users to make mistakes when working with data tables. If they request an operation that causes problems severe enough that the operation needs to be cancelled, the package raises an exception that informs them why it cannot do what they asked. If the operation can be completed but may cause issues for the user, the package issues a warning to alert them of the potential problem.

The package uses the standard Python methods for raising exceptions and issuing warnings. However, the standard way that Python prints warnings and exceptions, with the full stack trace and associated information, can be intimidating for users without a computer programming background. To make these messages easier to decipher, our package defines custom `sys.excepthook` and `warnings.showwarning` functions so that whenever the package raises an exception or issues a warning, it will be printed in a concise, approachable format for users. Warnings and exceptions from outside of the package are printed in the normal fashion. However, in an IPython notebook environment, we cannot control how warnings and exceptions are displayed, so this feature is a specific enhancement for users accessing the package through the command line or scripts..

## Results

To promote reproducibility, transparency and collaboration in the analysis of CPTAC proteogenomic data, we created a data API that explicitly links data access and data analysis. The API gives access to the CPTAC data from six cancer types: breast, colon, serous ovarian, endometrial, clear cell renal cell carcinoma, lung adenocarcinoma. As new datasets are published, they will become publicly available through the API. Data for each cancer type contains information on DNA, RNA, proteins, and clinical information (Figure 1). DNA data is derived from whole genome and whole exome sequencing of tumor and blood normal, and is processed to yield somatic mutation calls and somatic copy number variation calls. RNA data is from RNA-seq and is processed to yield transcriptomics and frequently circular RNA and micro-RNA. Protein data is from mass spectrometry based proteomics and contains global proteomics and phosphoproteomics. Clinical data contains patient demographic information, descriptive data for the tumor, and patient follow up. See the original publication for a detailed description of data acquisition and analysis methods^1–5^.

The data API is designed to provide frictionless access to quantitative data tables; its goal is to provide the data in the most convenient format with the least effort. The focus on quantitative data tables and not raw instrument data has several important benefits. First, these relatively small sized files do not necessitate a voluminous server nor special considerations for downloading, which enables the API to work on standard computers with standard internet connectivity. Second, these processed tables are also completely publicly available and do not contain any private germline information, further facilitating data access. Third, quantitative tables can be provided directly within the programming environment as a simple matrix variable, meaning that a user’s first interaction with data is as a properly parsed and loaded Python/Pandas dataframe (Figure 2A). In addition to single table access, the API also has native join functions which will merge multi-omics data types (Figure 2B). This simple access dramatically improves a user’s ability for interaction, exploration, and visualization (Figure 2C). Fourth, quantitative data files are the starting point of hypothesis testing and data analysis. Providing direct access to these data improves the transparency and reproducibility by ensuring all analysts are accessing the exact same data. This type of universal synchronization is not typically seen when analysts keep their own local version of data files, which are frequently altered and re-saved as re-normalized or filtered versions of the original. Moreover, independent file versions often lack detailed provenance of data manipulation. However, with an API, all data access is within a programming environment and any data manipulation is directly exposed in code following the data access. Finally, the data API represents a dramatically simpler method for users to access and analyze data. All data for the CPTAC cohorts is available in the same format, from the same simple software interface. There is no need to visit multiple repositories, e.g. sequencing and mass spectrometry data at multiple NCI Data Commons.

**Figure 2.**
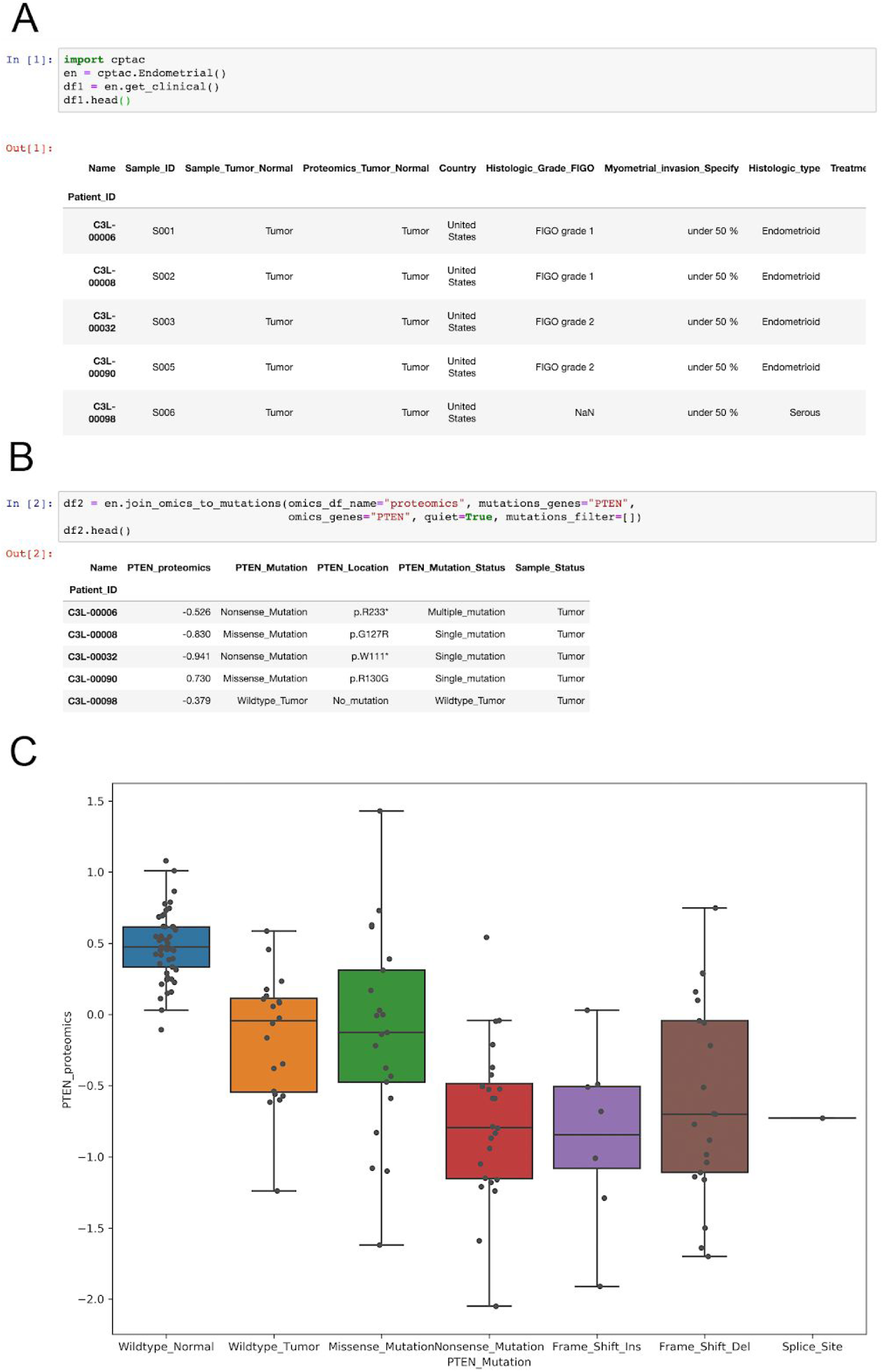
Getting data from the cptac API. A - The data API makes accessing CPTAC data simple, and returns data in a native Pandas dataframe. B - Merging different data types is facilitated by a suite of ‘join’ functions in the API. C - the joined mutation and proteomics data from panel B is shown with a boxplot from the `seaborn` Python graphing module. Example drawn from use case 2 in the cptac documentation (https://github.com/PayneLab/cptac/blob/master/docs/usecase2_clinical_attributes.ipynb).

### Versions

The data API is designed to keep track of formal versions of data. This is accomplished through a strict separation of the software code to load the data, and the actual data files. As with many large-scale projects, the data tables used in CPTAC analyses are periodically updated. Such adjustments are common in data analysis and using a formal data version helps to properly track changes. These updates are often prompted by new patient survival or treatment information from follow up visits. Distributing these updates throughout the consortium in a coordinated manner is best done through an official data release. Other reasons that prompt a data release include when the consortium has refined a software pipeline involved in generating the quantitative data tables (e.g. a different algorithm for processing raw sequencing or mass spectrometry data). The end-user can access these different versions when loading data into their Python environment (see Methods). When working with the data API, the default version is the current published version; however, a user can specify a different version.

### Tutorials and user support

One of the most important goals of the `cptac` package is to promote re-analysis of the consortium’s datasets. To facilitate this, we have created a comprehensive set of tutorials and use cases (Table 1). The tutorials cover the basic mechanics of accessing data through the API, and manipulating and merging data tables. The use cases are short examples that use the package to investigate real, biologically meaningful questions; they demonstrate how to use the API for hypothesis driven biological discovery. As the CPTAC datasets contain a variety of diverse data types, use cases often explore scientific questions that utilize integrated multi-omics data.

**Table 1:** User documentation. A list of tutorials and use cases to help users explore the data API, available at https://github.com/PayneLab/cptac/tree/master/docs. (a) The GSEApy module is available at https://gseapy.readthedocs.io/en/latest/introduction.html. (b) The BlackSheep module is available at https://blacksheep.readthedocs.io/en/master/.

|  |  |
| --- | --- |
| Tutorial 1: Data intro | Goes over the basics of how to install the package and access data |
| Tutorial 2: Pandas | More in-depth description of how to work with the tables using Pandas |
| Tutorial 3: Joining dataframes | Shows how to use built-in functions from cptac for joining tables of different data types |
| Tutorial 4: MultiIndex | Some tables provided by the package use multi-level column indexes in cases where multiple keys are required to uniquely identify each column. This tutorial describes unique aspects of working with these tables. |
| Tutorial 5: Updates | An explanation of how to access and work with data version updates, and package version updates. |
| Tutorial 6: Python and R | How access the Python API within R |
| Use case 1: Multi-omic integration | Data access and integration for multiple omics data types |
| Use case 2: Clinical covariates | Explores meta-data for correlation between clinical attributes |
| Use case 3: Clinical and acetylation | Compares acetylation levels between tumor subtypes |
| Use case 4: Mutations and omics | Studies the effects of DNA mutations on protein abundance |
| Use case 5: Enrichment analysis | Uses the GSEAPy<sup>a</sup> module to find enriched pathways |
| Use case 6: Derived molecular | Identifies correlation between proteomics and attributes derived from molecular data, e.g. MSI status |
| Use case 7: Trans genetic effect | Studies the effect of DNA mutations on the expression of a different protein |
| Use case 8: Outliers | Uses the Blacksheep<sup>b</sup> module to study outliers in expression values |
| Use case 9: Clinical outcomes | Uses patient follow up data to look for correlations between clinical and molecular data, and patient survival |
| Use case 10: Pathway overlay | Integrates quantitative molecular data with Reactome pathway maps |

The tutorials and use cases are written in interactive Python notebooks (Jupyter Notebooks). This format allows explanatory text, code, and output to be seamlessly integrated into a single document. The notebooks can be viewed on GitHub in the docs folder of the project repository (https://github.com/PayneLab/cptac/tree/master/docs). Users can also access interactive versions of the notebooks hosted on Binder (https://mybinder.org/v2/gh/PayneLab/cptac/master).In addition to the Python-based notebooks, tutorial 6 demonstrates how to access the API within the R programming environment.

Along with our tutorials and use cases, we also provide user support through our GitHub issues page. This page allows users to submit bug reports, feature requests, and other feedback about our package. Past questions and answers are publicly available for others to view for reference.

### Integrating with external bioinformatics tools

To enhance usability and functionality, the data API connects with several bioinformatics tools. The first category of tools are those that have a Python implementation. Working with these third-party packages is facilitated by our use of dataframes to encapsulate omics data, as many Python packages are written for Pandas. Gene set analysis is a frequent step in omics data analysis to find common biological functions among a specified set of genes. One of the first of these tools was GSEA^19^, which has been re-implemented in Python in the `gseapy` module (https://pypi.org/project/gseapy/). A tutorial demonstrating the use of `gseapy` is available in the `cptac` documentation. Another common task for CPTAC data is survival analysis. Several Python packages implement analyses like the Kaplan-Meier curves or the Cox proportional hazard test. These tests identify whether time to an event is affected by a specified variable, e.g. protein abundance, tumor grade, etc (Figure 3). The `cptac` tutorials demonstrate the `lifelines` package (https://pypi.org/project/lifelines/) using data from the API.

**Figure 3.**
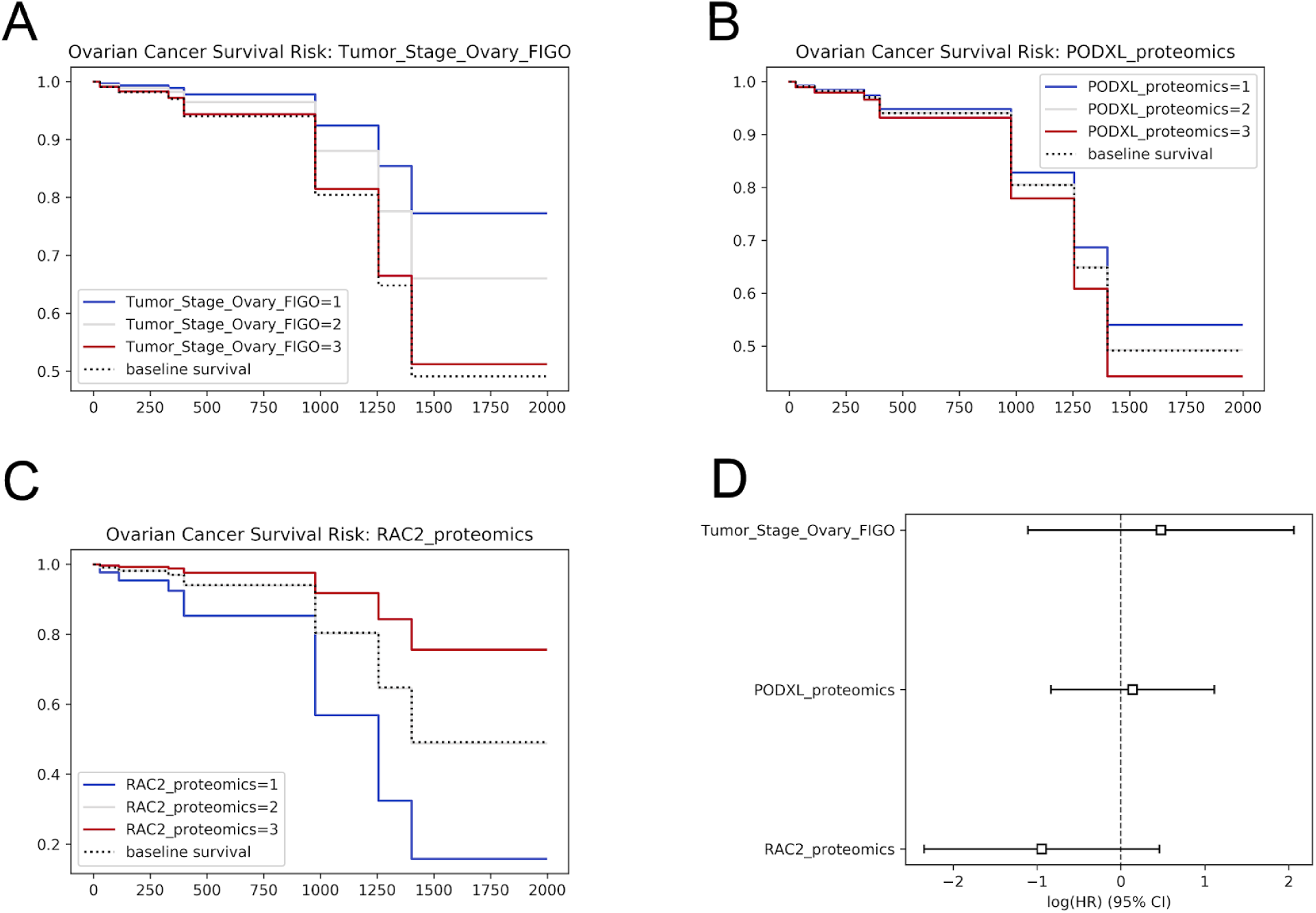
Survival analysis with CPTAC data. Using the `lifelines` Python module, we identify variables that impact patient survival in ovarian cancer. A Kaplan-Meier curve is shown to separate time to death based on: A - FIGO stage, B - protein expression of PODXL, C - RAC2. A Cox proportional hazard assessment is shown in D. Images generated using code in use case 9 of the `cptac` documentation (https://github.com/PayneLab/cptac/blob/master/docs/usecase9_clinical_outcomes.ipynb).

A second type of external tool that can be integrated with the data API are bioinformatics databases. Many large databases like Uniprot have web-services that allow for programmatic access to specific information. To integrate our CPTAC data with these, the API wraps REST calls to the database. For example, the API can give users a list of interacting proteins from both Uniprot and STRING by wrapping a call to their respective web-service. Some other databases either do not have a convenient REST-API or have a small enough database that it can be loaded in full. Proteins belonging to biological pathways are curated by Wikipathways. We have downloaded this open source information and integrated it into the data API, which users can access through simple function calls.

## Conclusions

Frictionless data access has become an important goal for science, as it promotes collaboration, transparency, reproducibility and data re-use. In the past decade, attitudes and expectations in the scientific community have changed with respect to data sharing. As data re-use extends the value of the financial investment beyond the original grant holder, various funding agencies have begun to set benchmarks for program success based on these more inclusive definitions of impact - as opposed to simply citation metrics. Although the sharing of raw data has a robust infrastructure for many primary data types, processed data tables are still frequently not shared in a simple and convenient manner. Here we present a data API for cancer proteogenomic data associated with the CPTAC consortium. By adopting the goals of the Data-as-a-Service model, our API radically improves access to these data.

As a generalized adaptation of the Data-as-a-Service model, our API’s guiding philosophy is that datasets should be accessible within a programming environment. Previous work with a similar on-demand goal is often achieved with a REST-API, which frequently returns data in a structured JSON format. Unfortunately, a JSON object requires parsing and reshaping to obtain the practical data matrix object. As a data matrix is the most common datatype for many scientific domains, our API directly gives users a native dataframe object. A second advantage of our data API implementation over a REST-API is that it obviates the need to host a webserver, making it simpler for creation and long-term maintenance. Numerous scientific projects, large and small, could improve the dissemination of their data through this mechanism.

## Acknowledgments

The authors acknowledge important testing and feedback on the API provided by members of the Payne lab and the Fenyö and Ruggles labs (New York University). The authors also thank Jason McDermott (Pacific Northwest National Laboratory) for initial conversations about the concept. This work was supported by the National Cancer Institute (NCI) CPTAC award [U24 CA210972], and by the Simmons Center for Cancer Research.

